# Towards standard practices for sharing computer code and programs in neuroscience

**DOI:** 10.1101/045104

**Authors:** Stephen J. Eglen, Ben Marwick, Yaroslav O. Halchenko, Michael Hanke, Shoaib Sufi, Padraig Gleeson, R. Angus Silver, Andrew P. Davison, Linda Lanyon, Mathew Abrams, Thomas Wachtler, David J. Willshaw, Christophe Pouzat, Jean-Baptiste Poline

**Affiliations:** Cambridge Computational Biology Institute, Department of Applied Mathematics and Theoretical Physics, University of Cambridge, UK; Department of Anthropology, University of Washington, Seattle, WA 98195-3100 USA; Department of Psychological and Brain Sciences, Dartmouth College, Hanover, NH 03755 USA; Institute of Psychology II, Otto-von-Guericke-University Magdeburg, 39106 Magdeburg, Germany; Center for Behavioral Brain Sciences, 39106 Magdeburg, Germany; Software Sustainability Institute, University of Manchester, UK; Department of Neuroscience, Physiology and Pharmacology, University College London, UK; Unité de Neurosciences, Information et Complexité, CNRS, Gif sur Yvette, France; International Neuroinformatics Coordinating Faility, Karolinska Institutet, Stockholm, Sweden; Department of Biology II, Ludwig-Maximilians-Universität at München, Germany; Institute for Adaptive and Neural Computation, School of Informatics, University of Edinburgh, UK; MAP5 Paris-Descartes University and CNRS UMR 8145, 45 rue des Saints-Pères, 75006 Paris, France; Henry H. Wheeler, Jr. Brain Imaging Center, Helen Wills Neuroscience Institute, University of California, Berkeley, USA

## Background

Many areas of neuroscience are now critically dependent on computational tools to help understand the large volumes of data being created. Furthermore, computer models are increasingly being used to help predict and understand the function of the nervous system. Many of these computations are complex and usually cannot be concisely reported in the methods section of a scientific article. In a few areas there are widely used software packages for analysis (e.g., SPM, FSL, AFNI, FreeSurfer, Civet in neuroimaging) or simulation (e.g. NEURON, NEST, Brian). However, we often write new computer programs to solve specific problems in the course of our research. Some of these programs may be relatively small scripts that help analyze all of our data, and these rarely get described in papers. As authors, how best can we maximize the chances that other scientists can reproduce our computations, find errors, or reuse our methods on their data? Is our research reproducible^1^?

To date, the sharing of computer programs underlying neuroscience research has been the exception (see below for some examples), rather than the rule. However, there are many potential benefits to sharing these programs, including increased understanding and reuse of your work. Furthermore, open source programs can be scrutinized and improved, whereas the functioning of closed source programs remains forever unclear^2^. Funding agencies, research institutes and publishers are all gradually developing policies to reduce the withholding of computer programs relating to research^3^. The Nature family of journals has published opinion pieces in favor of sharing whatever code is available, in whatever form^4,5^. Since October 2014, all Nature journals require papers to include a statement declaring *whether* the programs underlying central results in a paper are available. In April 2015 *Nature Biotechnology* offered recommendations for providing code with papers and began asking referees to give feedback on their ability to test code that accompanies submitted manuscripts^6^. In July 2015 F1000Research stated that "Software papers describing non-open software, code and/or web tools will be rejected" (http://f1000research.com/channels/f1000-faculty-reviews/for-authors/article-guidelines/software-tool-articles). Also in July 2015, BioMed Central introduced a minimum standards of reporting checklist for BMC Neuroscience and several other journals, requiring submissions to include a code availability statement and for code to be cited using a DOI or similar unique identifier^7^. We believe that all journals should adopt policies that highly encourage, or even mandate, the sharing of software relating to journal publications as this is the only practical way to check the validity of the work.

### What should be shared?

It may not be obvious what to share, especially for complex projects with many collaborators. As advocated by Claerbout and Donoho, for computational sciences the scholarship is not the article; the ”scholarship is the complete software […]^”8,9^. So, ideally, we should share all code and data needed to allow others to reproduce our work, but this may not be possible or practical. However, it is expected that the key parts of the work should be shared, e.g. implementations of novel algorithms or analyses. At a minimum, we suggest following the recommendation of submission of work to ModelDB^10^, i.e. to share enough code, data and documentation to allow at least one key figure from your manuscript to be reproduced. However, by adopting appropriate software tools, as mentioned in the next section, it is now relatively straightforward to share the materials required to regenerate *all* figures and tables. Code that already exists, is well tested and documented, and is reused in the analysis should be cited. Ideally, all other code should be communicated, including code that performs simple preprocessing or statistical tests, or code that deals with local computing issues such as hardware and software configurations. While this code may not be reusable, it will help others understand how analyses are performed, find potential mistakes, and aid reproducibility. Finally, if the work is computationally intensive and requires a long time to run (e.g. many weeks), one may prefer to provide a small "toy" example to demonstrate the code.

By getting into the habit of sharing as much as possible, not only do we help others who wish to reproduce our work (which is a basic tenet of the scientific method), we will be helping other members of our laboratory, or even ourselves in the future. By sharing our code publicly, we are more likely to write higher-quality code^11^, and we will know where to find it after we have moved on from the project^12^, rather than the code disappearing on a colleague’s laptop when they leave your group, or suffer some misfortune^13^. We also will be part of a community and benefit from the code shared by others, thus reducing software development time for ourselves and others.

### Simple steps to help you share code

Once you have decided *what* to share, here are some simple guidelines for *how* to share the work. Ideally, these principles should be followed throughout the lifetime of the research project, not just at the end when we wish to publish our results. Guidelines similar to these have been proposed in many areas of science^14–16^, suggesting that they are part of norms that are emerging across disciplines. In the ‘further reading’ section below, we list some specific proposals from other fields that expand on the guidelines we suggest here.

#### Version control

Use a version control system (such as Git) to develop the code^17^. The version control repository can then be easily and freely shared with others using sites such as http://github.com^18^ or https://bitbucket.org. These sites allow you fine control over private versus public access to your code. This means that you can keep your code repository private during its development, and then publicly share the repository at a later stage e.g. at the time of publication, although we recommend opening the code from the start of the project. It also makes it easy for others to contribute to your code, and to adapt it for their own uses.

#### Persistent URLs

Generate stable URLs (such as a DOI) for key versions of your software. Unique identifiers are a key element in demonstrating the integrity and reproducibility of research^19^, and allow referencing of the exact version of your code used to produce figures. DOIs can be obtained freely and routinely with sites such as http://zenodo.org and http://figshare.com. If your work includes computer models of neural systems, you may wish to consider depositing these models in established repositories such as ModelDB^10^, Open Source Brain^20^ or NITRC^21^. Some of these sites allow for private sharing of repositories with anonymous peer reviewers. Journal articles that include a persistent URL to code deposited in a trusted repository meet the requirements of level two of the ‘analytic methods (code) transparency’ standard of the TOP guidelines^14^.

#### License

Choose a suitable license for your code to assert how you wish others to reuse your code. For example, to maximize reuse, you may wish to use a permissive license such as MIT or BSD^22^. Licenses are also important to protect you from others misusing your code. Visit http://choosealicense.com/ to get a simple overview of which license to choose, or http://www.software.ac.uk/resources/guides/adopting-open-source-licence for a detailed guide.

#### Etiquette

When working with code written by others, observe Daniel Kahneman’s ‘reproducibility etiquette’^23^ and have a discussion with the authors of the code to give them a chance to fix bugs or respond to issues you have identified before you make any public statements. Cite their code in an appropriate fashion.

#### Documentation

Contrary to popular expectations, you do not need to write extensive documentation or a user’s guide for the code to still be useful to others^4^. However, it is worth providing a minimal README file to describe what the code does, and how to run it. For example, you should provide instructions on how to regenerate key results, or a particular figure from a paper. Literate programming methods, where code and narrative text are interwoven in the same document, make documentation semi-automatic and can save a lot of time when preparing code to accompany a publication^24,25^. However, these methods admittedly take more time to write in the first instance, and you should be prepared to rewrite documentation when rewriting code. In any cases, well-documented code allows for easier re-use and checking.

#### Tools

Consider using modern, widely used software tools that can help with making your computational research reproducible. Many of these tools have already been used in neuroscience and serve as good examples to follow, for example Org mode^26^, IPython/Jupyter^27^ and Knitr^28^. Virtualization environments, such as VirtualBox appliances and Docker containers, can also be used to encapsulate or preserve all of the computational environment so that other users can run your code without having to install numerous dependencies^29^.

#### Case studies

In addition to the examples listed above in Tools^26–28^, there are many prior examples to follow when sharing your code. For example, some prominent examples of reproducible research in computational neuroscience include Vogels et al.^30^ and Waskom et al.^31^; see https://github.com/WagnerLabPapers for details. The ModelDB repository contains over 1000 computational models deposited with in-structions for reproducing key figures to papers e.g. https://senselab.med.yale.edu/ModelDB/showModel.cshtml?model=93321 for a model of activity-dependent conductances^32^.

#### Data

Any experimental data collected alongside the software should also be released or made available. For small datasets, this could be stored alongside the software, although it may be preferable to store experimental data separately in an appropriate repository. Both PLOS and Scientific Data maintain useful lists of subject-specific and general repositories for data storage, see http://journals.plos.org/plosbiology/s/data-availability#loc-recommended-repositories and http://www.nature.com/sdata/data-policies/repositories.

#### Standards

Use of (community) standards, where appropriate, should be encouraged, in particular use of non-proprietary formats to enable long-term accessibility. In computational neuroscience for example, PyNN^33^ and NeuroML^34^ are widely used formats for making models more accessible and portable across multiple simulators. Neuroimaging data and results can be organized using BIDS^35^.

#### Tests

Testing the code has long been recognized as a critical step in the software industry but the practice is not widely adopted yet by researchers. We recommend including test suites that demonstrate the code is producing the correct results^36^. These tests can be at a low level (testing each individual function, called unit testing) or at a higher level (e.g. testing that the program yields correct answers on simulated data)^37^. With public data available, it is often straightforward to have a test verifying that published results can be recomputed. Linking tests to continuous integration services (such as Travis CI, https://travis-ci.org) allows these tests to be automatically run each time a change is made to the code, ensuring failing tests are immediately flagged and can be dealt with quickly.

#### User support

Although some people are eager to provide support for their code after it has been published, others may feel that they do not want to be burdened by e.g. feature requests. One simple suggestion to avoid this is to establish a user community for the code^38^. This could be as simple as creating a mailing list or asking for issues to be posted on a github repository.

###### Further reading (note to editor: please make this a box feature)

Khodiyar, V. 2015. Code Sharing — read our tips and share your own. Scientific Data Blog, February 19, 2015. http://blogs.nature.com/scientificdata/2015/02/19/code-sharing-tips/

Kitzes, J., Turek, D., & Deniz, F. (Eds.). 2017. The Practice of Reproducible Research: Case Studies and Lessons from the Data-Intensive Sciences. Oakland, CA: University of California Press. https://www.practicereproducibleresearch.org/

Leveque, R. 2013. Top ten reasons to not share your code (and why you should anyway). SIAM News, April 2013, https://sinews.siam.org/Details-Page/top-ten-reasons-to-not-share-your-code-and-why-you-should-anyway

Stodden, V., M. McNutt, D. H. Bailey, E. Deelman, Y. Gil, B. Hanson, M. A. Heroux, J.P. A. Ioannidis and M. Taufer 2016. Enhancing reproducibility for computational methods. Science 354(6317):1240. DOI: http://doi.org/10.1126/science.aah6168

Stodden V., & Miguez, S., 2014. Best practices for computational science: software infrastructure and environments for reproducible and extensible research. Journal of Open Research Software. 2(1), p.e21. DOI: http://doi.org/10.5334/jors.ay

Stodden, V., Leisch, F., & Peng, R. (Eds.). 2014. Implementing reproducible research. CRC press, Chapman and Hall. https://osf.io/s9tya/

Halchenko, Y. O. and Hanke, M. 2015. Four aspects to make science open "by design" and not as an after-thought. GigaScience, 4. DOI: http://doi.org/10.1186/s13742-015-0072-7

Sandve, G. K., Nekrutenko, A., Taylor, J., & Hovig E 2013. Ten simple rules for reproducible computational research. PLoS Comput Biol 9:e1003285.

###### StackExchange and related projects

StackExchange is a network of free and highly active question-and-answer websites. Two members of the network are relevant to questions of code sharing: http://stackoverflow.com which is dedicated to questions about programming in any language in any context, and http://academia.stackexchange.com/questions/tagged/reproducible-research which is focused questions relating to reproducible research in academic context. A related project is https://neurostars.org/ which is a similar free public Q&A website focused on neuroinformatics questions, and with many questions on software packages, etc.

###### Scientists for Reproducible Research

This is an international multi-disciplinary email list that discusses a wide range of issues relating to code sharing: https://groups.google.com/forum/#!forum/reproducible-research

###### GitHub

GitHub is an online repository for computer code and programs that has a large community of researchers that develop and share their code openly on the site. GitHub is the largest and most active code sharing site (others include BitBucket and GitLab) and has convenient tools for facilitating efficient collaborative coding^39,40^. If you are using an open source program you may find a community of users and developers active on GitHub, where you can ask questions and report problems.

### Closing remarks

Changing the behaviors of neuroscientists so that they make their code more available will likely be resisted by those who do not see the community benefits as outweighing the personal costs of the time and effort required to share code^41^. The community benefits, in our view, are obvious and substantial: we can demonstrate more robustly and transparently the reliability of our results, we can more easily adapt methods developed by others to our data, and the impact of our work increases as others can similarly reuse our methods on their data. Thus, we will endeavor to lead by example, and follow all these practices as part of our future work in all scientific publications. Even if the code we produce today will not run ten years from now, it will still be a more precise and complete expression of our analysis than the text of the methods section in our paper.

However, exhortations such as this article are only a small part of making code sharing a normal part of doing neuroscience; many other activities are important. All researchers should be trained in sound coding principles; such training is provided by organizations such as Software Carpentry^37^ or Data Carpentry and through national neuroinformatics initiatives, e.g. http://python.g-node.org. Furthermore, we should request code and data when reviewing, and submit to and review for journals that support code sharing. Grant proposals should be checked for mentions of code availability, and we should encourage efforts toward openness in hiring, promotion, and reference letters^42^. Funding agencies and editors should also consider mandating code sharing by default. This combination of efforts on a variety of fronts will increase the visibility of research accompanied by open source code, and demonstrate to others in the discipline that code sharing is a desirable activity that helps move the field forward.

We believe that the sociological barriers to code sharing are harder to overcome than the technical ones. Currently, academic success is strongly linked to publications and there is little recognition for producing and sharing code. Code may also be seen as providing a private competitive advantage to researchers. We challenge this view and propose that code be regarded as part of the research products and part of the publication in which should be shared by default, and that there should be an obligation to share code for those conducting publicly funded research. We hope the code availability review (*CITE JOURNAL EDITORIAL HERE*) will help establish such sharing as the norm. Moreover, we are advocating for code sharing as part of a broader culture change embracing transparency, reproducibility, and re-usability of research products.

## Acknowledgments

This article is based upon discussions from a workshop to encourage sharing in neuroscience, held in Cambridge, December 2014. It was financially supported and organized by the International Neuroinformatics Coordinating Facility (http://www.incf.org), with additional support from the Software Sustainability institute (http://www.software.ac.uk). Michael Hanke was supported by funds from the German federal state of Saxony-Anhalt and the European Regional Development Fund (ERDF), Project: Center for Behavioral Brain Sciences.

